# Integrative pharmacogenomics to infer large-scale drug taxonomy

**DOI:** 10.1101/046219

**Authors:** Nehme El-Hachem, Deena M.A. Gendoo, Laleh Soltan Ghoraie, Zhaleh Safikhani, Petr Smirnov, Christina Chung, Kenan Deng, Ailsa Fang, Erin Birkwood, Chantal Ho, Ruth Isserlin, Gary D. Bader, Anna Goldenberg, Benjamin Haibe-Kains

## Abstract

Identification of drug targets and mechanism of action (MoA) for new and uncharacterized drugs is important for optimization of drug efficacy. Current MoA prediction approaches largely rely on prior information including side effects, therapeutic indication and/or chemo-informatics. Such information is not transferable or applicable for newly identified, previously uncharacterized small molecules. Therefore, a shift in the paradigm of MoA predictions is necessary towards development of unbiased approaches that can elucidate drug relationships and efficiently classify new compounds with basic input data. We propose a new integrative computational pharmacogenomic approach, referred to as Drug Network Fusion (DNF), to infer scalable drug taxonomies that relies only on basic drug characteristics towards elucidating drug-drug relationships. DNF is the first framework to integrate drug structural information, high-throughput drug perturbation and drug sensitivity profiles, enabling drug classification of new experimental compounds with minimal prior information. We demonstrate that the DNF taxonomy succeeds in identifying pertinent and novel drug-drug relationships, making it suitable for investigating experimental drugs with potential new targets or MoA. We highlight how the scalability of DNF facilitates identification of key drug relationships across different drug categories, and poses as a flexible tool for potential clinical applications in precision medicine. Our results support DNF as a valuable resource to the cancer research community by providing new hypotheses on the compound MoA and potential insights for drug repurposlng.

## INTRODUCTION

Continuous growth and ongoing deployment of large-scale pharmacogenomic datasets has opened new avenues of research for the prediction of biochemical interactions of small molecules with their respective targets and therapeutic effects, also referred to as drug mechanisms of action (MoA). Development of computational methods to predict MoA of new compounds is an active field of research in the past decade^1,2^. Despite major advancements in this field, key challenges still remain in the (*i*) design of classification approaches that rely on minimal drug characteristics to classify drugs, and (*ii*) selection and integration of complementary datasets to best characterize drugs’ effects on biological systems.

The notion of a ‘minimalist’ approach to represent similarities among drug compounds has been extensively explored, with varying results. Several computational strategies have solely relied on chemical structure similarity to infer drug-target interactions ^3–5^, based on the assumption that structurally-similar drugs share similar targets, and ultimately, similar pharmacological and biological activity ^6^. However, sole dependence on chemical structure information fails to consider drug-induced genomic and phenotypic perturbations, which directly connect with biological pathways and molecular disease mechanisms ^7,8^. Seminal approaches by Iorio et al. ^9^ and Iskar et al. ^10^ leveraged drug-induced transcriptional profiles from Connectivity Map (CMAP) ^11^ towards identification of drug-drug similarities and MoA solely based on gene expression profiles ^12^. The major limitation of CMAP however is the lack of global scope, as only 1,309 drugs are characterized across 5 cancer cell lines ^11^. Other methods have integrated prior knowledge such as adverse effects annotations ^13,14^ and recent approaches showed that integrating multiple layers of information had improved ATC prediction for FDA-approved drugs ^15^. While these initiatives have paved great strides towards characterizing drug MoA, comparing the consistency of such efforts towards prediction of new, uncharacterized small molecules remains a challenge.

Ongoing efforts towards understanding drug behavior have yet to capitalize on newly developed, high-throughput data types towards providing improved classification of drug action mechanisms. The published CMAP has recently been superseded by the L1000 dataset from the NIH Library of Integrated Network-based Cellular Signatures (LINCS) consortium ^16^, which has expanded upon the conceptual framework of CMAP and contains over 1.8 million gene expression profiles spanning 20,413 chemical and genetic perturbations. A recent study of the LINCS data showed that structural similarity are significantly associated with similar transcriptional changes ^8^. While the L1000 dataset provides an unprecedented compendium of both structural and transcriptomic drug data, its integration with other pharmacogenomics data types has not been explored extensively.

The advent of high-throughput *in vitro* drug screening promises to provide additional insights into drug MoA. The pioneering initiative of the NCI60 panel provided an assembly of tumour cell lines that have been treated against a panel of over 100,000 small molecules ^17,18^, and served as the first large-scale resource enabling identification of lineage-selective small molecule sensitivities ^19^. However, its relatively small number of cancer cell lines (n=59) restricted the relevance of these data for prediction of drug MoA. To address this issue, the Cancer Therapeutics Response Portal (CTRP) initiative screened 860 cancer cell lines against a set of 481 small molecule compounds ^20,21^, which makes it the largest repository of *in vitro*drug sensitivity measurements to date. Individual assessment of these *in vitro* sensitivity datasets have highlighted their use towards determining mechanism of growth inhibition, and inference of MoA of compounds from natural products. It remains to be demonstrated, however, whether integration of drug sensitivity data with other drug-related data, such as drug structures and drug-induced transcriptional signatures, can be used to systematically infer drug MoA.

To efficiently harness these recent high-throughput datasets, we have developed a scalable approach that maximizes complementarity between different data types to provide a complete landscape of drug-drug relationships and similarities. We leveraged our recent Similarity Network Fusion algorithm ^22^ to integrate drug structure, sensitivity, and perturbation data towards developing a large-scale molecular drug taxonomy, called Drug Network Fusion (DNF) (Fig. 1). We demonstrate how the resulting integrative drug taxonomies improve characterization of drug MoA compared to taxonomies based on single data types. We demonstrate how these new drug networks can be harnessed to evaluate drug-drug similarities across drug targets and ATC classification benchmarks. Importantly, we show how the DNF taxonomy can be used to infer MoA for new compounds that lack deep pharmacological and biochemical characterization.

**Figure 1.**
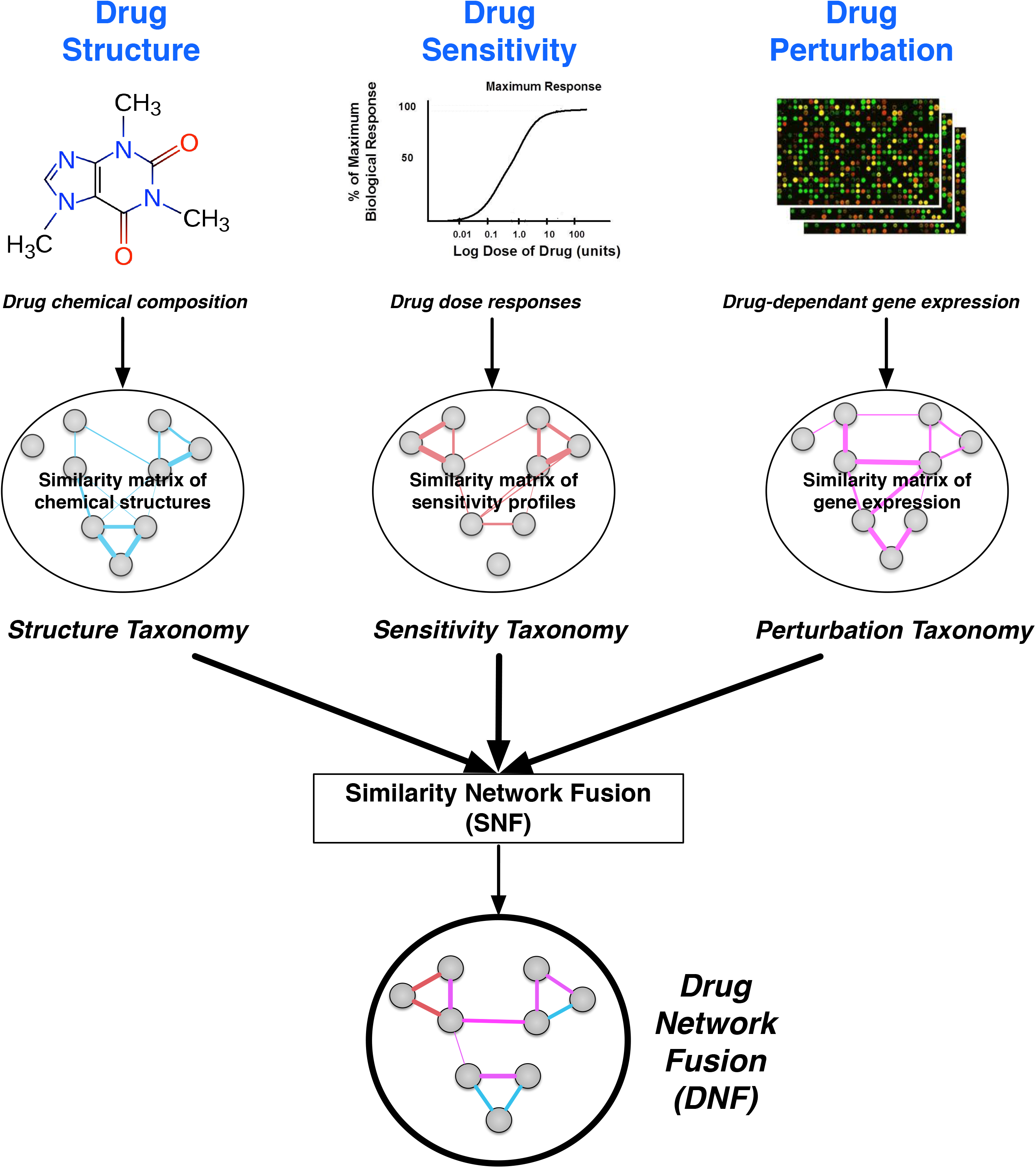
Schematic representation of the SNF method and its use towards integration of different types of drug information. Datasets representing drug similarity, drug sensitivity, and drug perturbation profiles are first converted into drug-drug similarity matrices. Similarity matrices are fully integrated within the SNF method to generate a large-scale, multi-tier, Drug Fusion Network (DNF) taxonomy of drug-drug relationships.

## RESULTS

We developed the DNF approach to generate a large-scale molecular taxonomy by integrating drug similarity matrices from structural information, transcriptomic perturbation, and sensitivity profiles **(Supplementary Fig. 1)**. Drug structure profiles (SMILES representations) were extracted from the PubChem database. Drug perturbation signatures, representing drug-induced gene expression changes, were extracted from the recent LINCS L1000 dataset. Drug sensitivity profiles representing cell line viability across cancer cell lines were extracted from our PharmacoGx platform ^23^, which contains pharmacological profiles of several hundred cell lines generated by the CTRP ^20^,^21^ and NCI60 ^19^initiatives. By integrating these three data types, we have generated a similarity network composed of over 200 drugs **(Supplementary Fig. 2)**. The drug similarity matrices computed from single data layers and fusion estimates are provided in **Supplementary Tables 1 and 2** for the CTRPv2 and NCI60 taxonomies.

To assess the relevance and benefits of our integrative drug taxonomies we first tested whether the different data layers were redundant or complementary. We then tested whether DNF enables prediction of drug MoA. While definitions may vary in the literature, for the context of this study, we refer to drug mechanisms of action specifically as determining drug targets for query drugs, as well as identification of the anatomical therapeutic classification (ATC) for different drugs. We further tested whether DNF enables clustering of drugs sharing common action mechanisms, and we demonstrated how identified drug communities from this clustering may be used towards novel discoveries for drug repositioning approaches and clinical applications.

### Complementarity of drug structure, perturbation, and sensitivity profiles

We investigated the potential complementarity between drug structure, perturbation, and sensitivity profiles to assess their potential for drug taxonomies as part of DNF. We generated similarity networks (similarity matrices) representing each of the single-layer drug taxonomies of drug structure, drug perturbation, and drug sensitivity (**Supplementary Tables 1 and 2**). In brief, we used the Tanimoto index to calculate the similarity between two drug structures. We used Pearson correlation to quantify the extent to which pairs of drugs affect similar genes at the expression level or whether drugs kill the same cancer cell lines for drug perturbation and drug sensitivity similarity matrices, respectively. We then computed the correlation between all pairs of similarity matrices (Fig. 2, **Supplementary Table 3**) using the spearman correlation. The single-layers are only weakly correlated, with a maximum absolute spearman correlation coefficient of 0.091 between the perturbation and sensitivity layers using the CTRPv2 sensitivity dataset (Fig. 2A), as well as a maximum of 0.085 between the structure and sensitivity layers using the NCI60 sensitivity dataset (Fig. 2B). In contrast, the integrative drug taxonomy (DNF) is more highly correlated to each of the single layers, across both CTRP (Fig. 2A) and NCI60 (Fig. 2B). These findings highlight that the structural, sensitivity, and transcriptional perturbation data are non-redundant, thereby presenting a diversity of drug-drug relationships across the set of drugs that commonly share all three data types. These findings also highlight that the DNF network maximizes fusion between the three data types, such that it is not driven by only a single layer.

**Figure 2.**
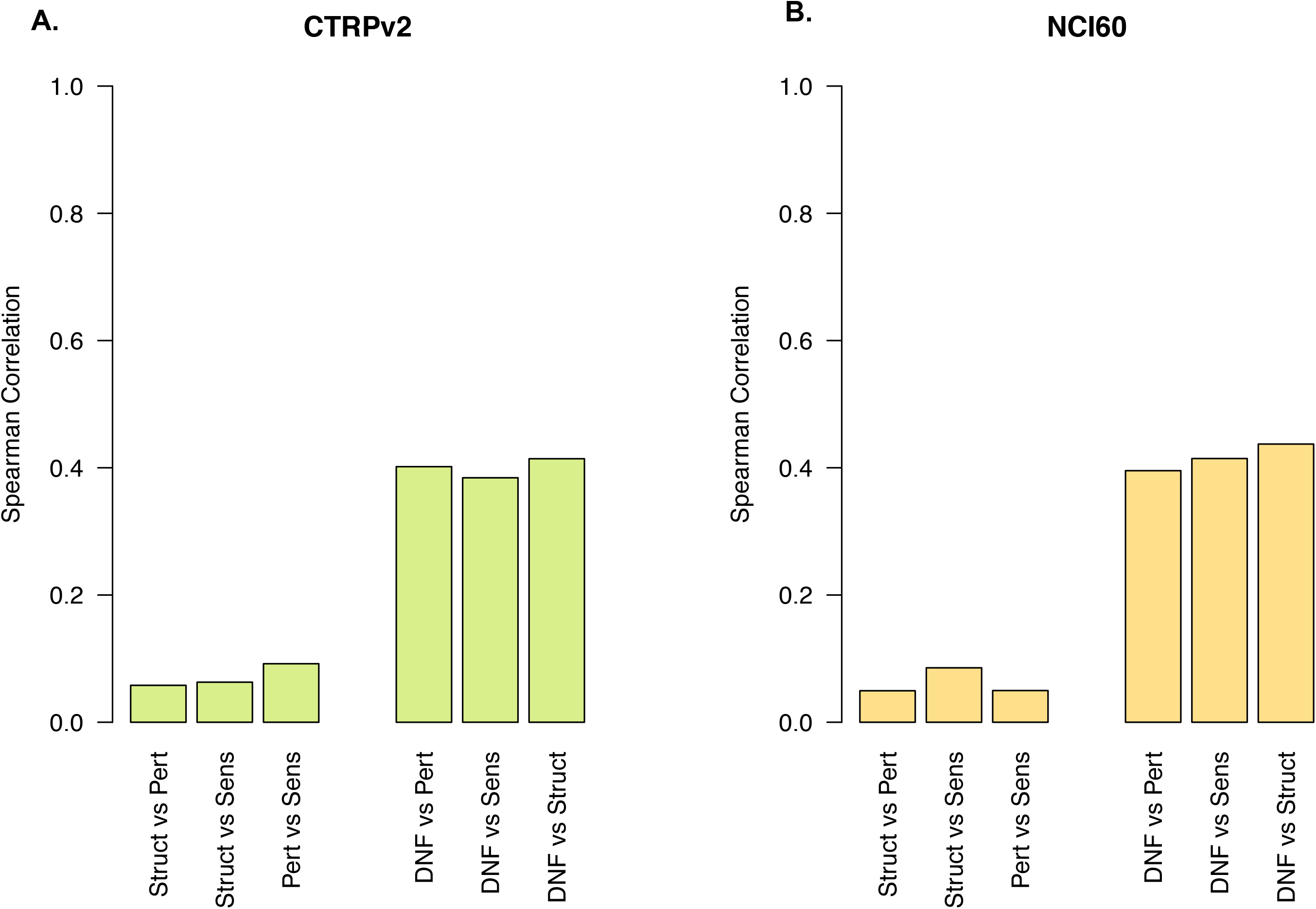
Complementarity of drug information across drug taxonomies. Spearman correlation between all pairs of single-layer similarity matrices (drug structure, drug perturbation, drug sensitivity) are depicted. Correlations between the integrative drug taxonomy (DNF) and each of the single-layer similarity matrices are also show. Data are shown for both (**A**) drug taxonomy using CTRPv2 and (**B**) drug taxonomy using the NCI60 sensitivity datasets.

### Performance of drug taxonomies against known drug targets

Determining novel drug-target interactions opens new avenues for drug repurposing efforts, and suggests mechanisms by which drugs can operate in cells. We explored the relevance of our DNF taxonomy by demonstrating its predictive value towards identification of drug targets. Of the 239 drugs represented in our DNF taxonomy generated using CTRPv2, 141 could be matched against the drug target benchmarks **(Supplementary Fig. 2)**. Similarity, for the DNF taxonomy generated using the NCI60 dataset, 86 drugs out of 238 drugs could be matched to known drug target **(Supplementary Fig. 2)**. We assessed the predictive value of our single-data layer and integrative drug taxonomies (DNF) against known drug targets **(Supplementary Fig. 3)**. We performed a receiver operating characteristics (ROC) analysis to quantify how well our drug taxonomies align with established drug target, by statistically comparing the area under the ROC curve (AUROC) values for each drug taxonomy under evaluation ^24^. Similarly, we calculated the area under the precision-recall (PR) curve (AUPRC)25. Of the three single-layer taxonomies validated against annotated drug targets from CTRPv2, the drug sensitivity layer outperformed the structure and perturbation taxonomies (AUROC of 0.83, 0.71 and 0.64 for sensitivity, structural and perturbation data layers, respectively) (Fig. 3A). Importantly, DNF yielded the best predictive value (AUROC of 0.89), and was significantly higher than any single-layer taxonomy (one-sided t test p-value < 1E-16, **Supplementary Table 4**). We further computed PR curves (Fig. 3B). These curves demonstrate that DNF supersedes the single layers taxonomies except for sensitivity, where it performs equivalently (AUPRC of 0.413 and 0.406 for DNF and drug sensitivity, respectively; Fig. 3B).

**Figure 3.**
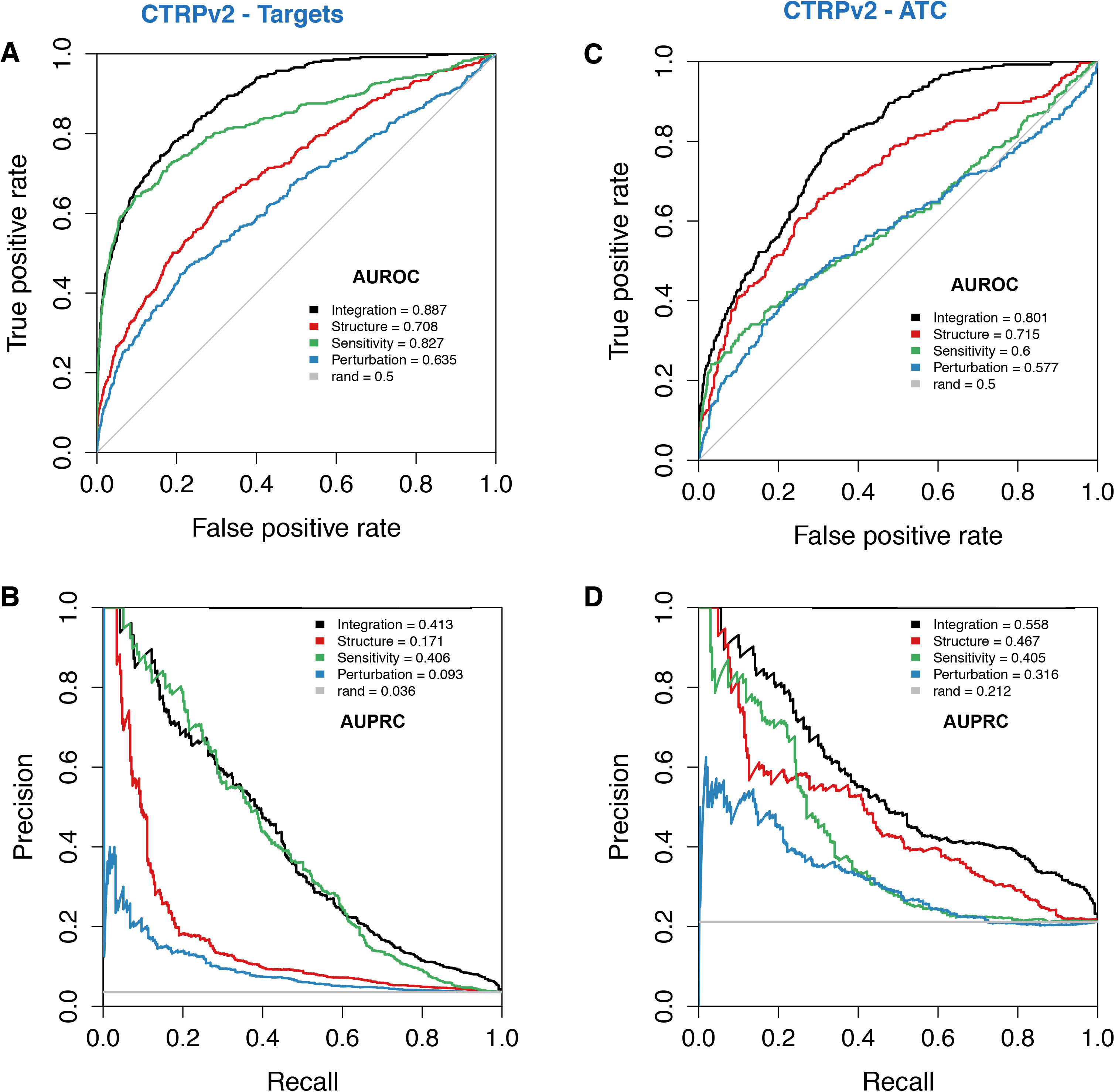
Validation of the DNF taxonomy using CTRPv2 sensitivity data and single dataset taxonomies against the ATC and Drug-target benchmarks. ROC and PR curves are shown for each of the taxonomies generated with the CTRPv2 sensitivity dataset, tested against ATC annotations and drug-target information from CHEMBL or internal benchmarks. Lines representing random (“rand”) classifications are drawn in grey for both ROC and PR curves. (**A**) ROC curve against drug-targets (**B**) PR curve against drug-targets (**C**) ROC curve against ATC drug classifications (**D**) PR curve against ATC drug classifications

We further tested the predictive value of our DNF taxonomy using the set of drug sensitivity profiles obtained from the NCI60 dataset, where a much smaller panel of cell lines has been screened (60 vs 860 cell lines for NCI60 and CTRPv2, respectively; Supplementary Fig. 2). Our evaluation of single-layer taxonomies demonstrates that drug similarities based on structure and sensitivity profiles were the most efficient in predicting drug-target associations (AUROC of 0.8 for both layers; Supplementary Fig. 4A) compared to perturbation (AUROC of 0.62; Supplementary Fig. 4A). DNF was significantly more predictive of drug-target associations compared to single-layer taxonomies (AUROC of 0.88 and one-sided superiority test p-values < 0.05, Supplementary Fig. 4A, **Supplementary Table 4** and AUPRC of 0.552; Supplementary Fig. 4B).

### Performance of drug taxonomies against anatomical classification (ATC)

Predicting the anatomical therapeutic chemical classification (ATC) of a drug provides insights about its pharmacological mechanism, and ultimately presents new potential indications for previously uncharacterized drugs. We demonstrated the relevance of our DNF network by testing its predictive value for ATC drug classifications (Fig. 3, Supplementary Fig. 4). We explored the value of the DNF taxonomies towards classifying drugs up to ATC level 4, which reports the chemical/therapeutic/pharmacological subgroup of a given drug ^26^. A total of 51 and 72 drugs could be matched against the ATC benchmarks for the CTRPv2 and NCI60 taxonomies, respectively. We implemented a similar benchmarking approach to that previously conducted for drug target classification. We observed that drug sensitivity was no longer the most predictive layer for ATC classification, and instead exhibited comparable predictive power to drug perturbation (Fig. 3C, Supplementary Fig. 4C). Conversely, the structure-based taxonomy (Fig. 3C, Supplementary Fig. 4C) was the most predictive amongst single-layer taxonomies (AUROC of 0.72, 0.6 and 0.58 for structure, sensitivity, and perturbation layers, respectively, for the CTRPv2 taxonomy; see Supplementary Fig. 4 for the NCI60 taxonomy). DNF significantly outperformed single-layer taxonomies (Fig. 3C-D, Supplementary Fig. 4C-D) (AUROC of 0.8 and 0.85 for DNF based on CTRPv2 and NCI60, respectively, with one-sided t test p-value < 0.05; and AUPRC of 0.558 and 0.492 vs. random classifiers’ AUPRCs of 0.212 and 0.095, respectively) Fig. 3C, Supplementary Fig. 4C, **Supplementary Table 4**).

### Identification of drug communities

We assessed the biological relevance of DNF in discovering drugs with similar MoA by applying the affinity cluster propagation algorithm ^27^ to identify clusters of highly similar drugs, referred to as *drug communities.* These communities can be represented by their most representative (‘exemplar’) drug, and similarities between communities are subsequently represented a network where each node is labeled by the exemplar drug. Community detection was implemented on the full set of 239 and 238 drugs of the DNF taxonomy (based on using CTRPv2 and NCI60 sensitivity data; Fig. 4 and Supplementary Fig. 5 respectively).

**Figure 4.**
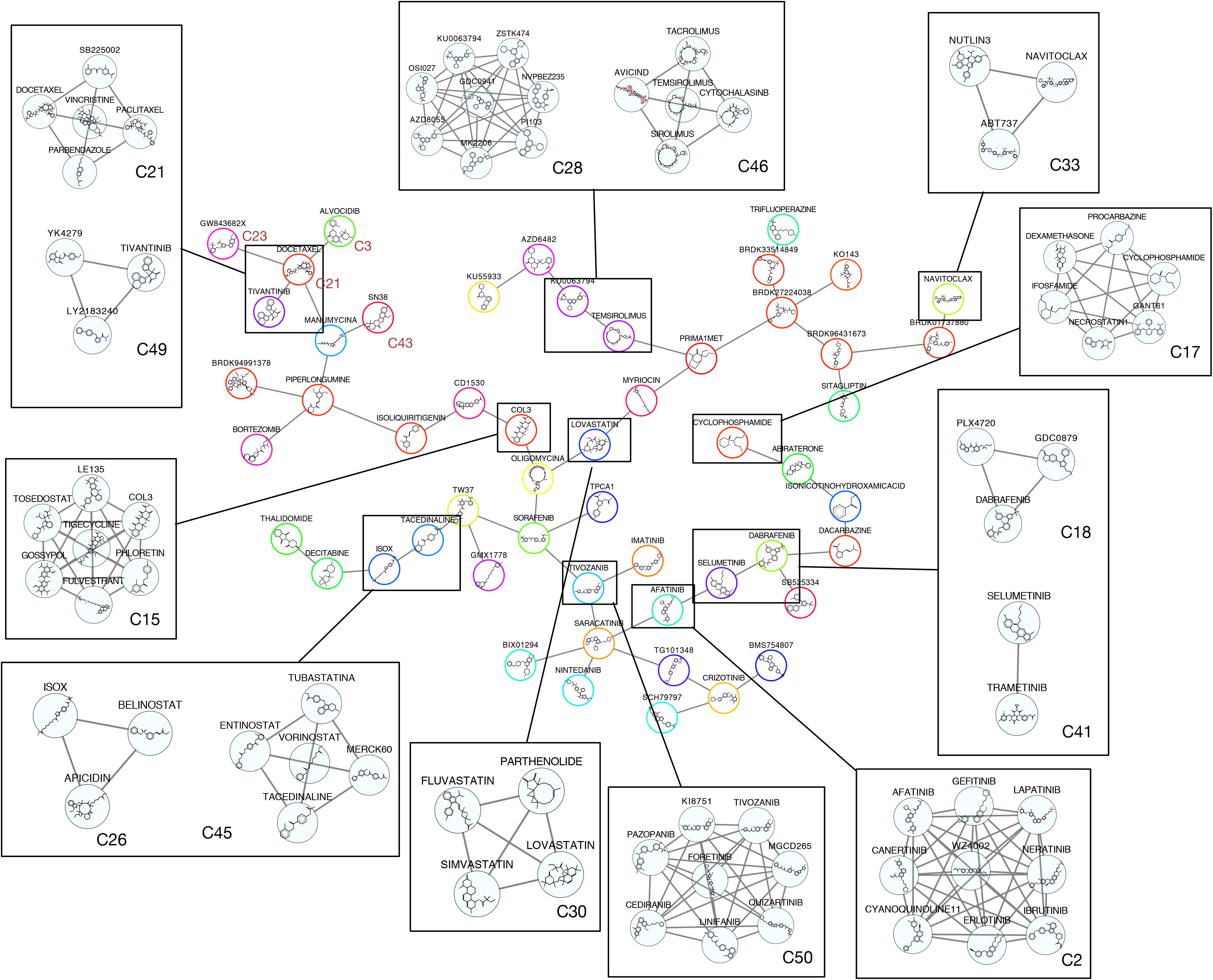
Network representation of 51 exemplar drugs that are representative of the drug communities identified by the DNF taxonomy using CTRPv2 sensitivity data. Each node represents the exemplar drugs, and node sizes reflect the size of the drug community represented by the exemplar node. Nodes are colored to reflect shared MoA as determined using known drug targets. Communities sharing similar MoA and proximity in the network are highlighted, with the community number indicated next to each community. Drug communities pertaining to the super-community are labelled in red.

We identified 53 communities in the CTRPv2 DNF (**Supplementary Table 5**), which resulted in a set of 39 drug communities with at least 2 drugs with known drug targets (permutation test p-value < 1E-4; **Supplementary Table 6**). Overall, DNF has produced a consistent classification of drugs for a variety of known functional classes (**Supplementary Table 7**). Our classifications recapitulate most of the protein target-drug associations represented ln CTRPv2 (Fig. 4). Receptor tyrosine kinases and non-receptor tyrosine kinases, including EGFR/ERBB2 (community C2), MEK1/2 (C41), TGFRB1 (C39), BRAF (C18), IGFR-1 (C6) KDR/FLT1 (C50) inhibitors. In addition to PI3K/mTOR inhibitors (C28), epigenetic regulators: HDACs (C45) and DNMT1 (C20) inhibitors, HMG CoA (C30) and proteasome inhibitors (C7) (**Supplementary Tables 5 and 8**). Notably, all BRAF (V600E mutation) inhibitors were classified correctly, which include drugs already tested in metastatic melanoma (community C18: dabrafenib, GDC0879, PLX4720; Fig. 4) and mitogen-activated protein kinase/ERK kinase (MEK) inhibitors (C41: namely trametinib and selumetinib; Fig. 4). BRAF regulates the highly conserved MAPK/ERK signaling pathway, and BRAF mutational status has been proposed as a biomarker of sensitivity towards selumetinib and other MEK inhibitors^28,29^. This explains the tight connection of these two communities (Fig. 4).

Using the NCI60 DNF, we identified 51 communities (Supplementary Fig. 5, **Supplementary Table 9**) with 20 of those containing at least two known drug targets (permutation test p-value < 1E-4; **Supplementary Table 10**). In this case, we also identified a number of well-characterized drug communities. These include the community composed of EGFR inhibitors (C20; Supplementary Fig. 5). The community C14, including cardiac glycosides also concur with the study of Khan et al. ^7^, showing that these compounds inhibit Na+/K+ pumps in cells. Bisacodyl, a laxative drug, is part of the C14 community, which concurs with Khan et al. who demonstrated a sharing a MoA similar to cardiac glycosides, despite its structural dissimilarity to that class of compounds ^7^.

### Enrichment of DNF drug communities for drug targets and ATC classifications

We conducted a quantitative community enrichment analysis to test whether DNF succeeds in identifying drug communities that are enriched for drug targets and ATC classifications. A fisher test was conducted between all the drugs in each community versus all drugs attributed to a specific drug target or ATC class. This approach allowed us to test which specific drug targets or ATC classifications are significantly enriched in the computed communities (Fig. 5, Supplementary Fig. 6), independent of the benchmarking analyses. The analysis was conducted using all of the communities of DNF based on CTRPv2 (n=53 communities, Fig. 5, **Supplementary Table 11**) as well as DNF based on NCI60 sensitivity data (n=51 communities, Supplementary Fig. 6, **Supplementary Table 12**).

**Figure 5.**
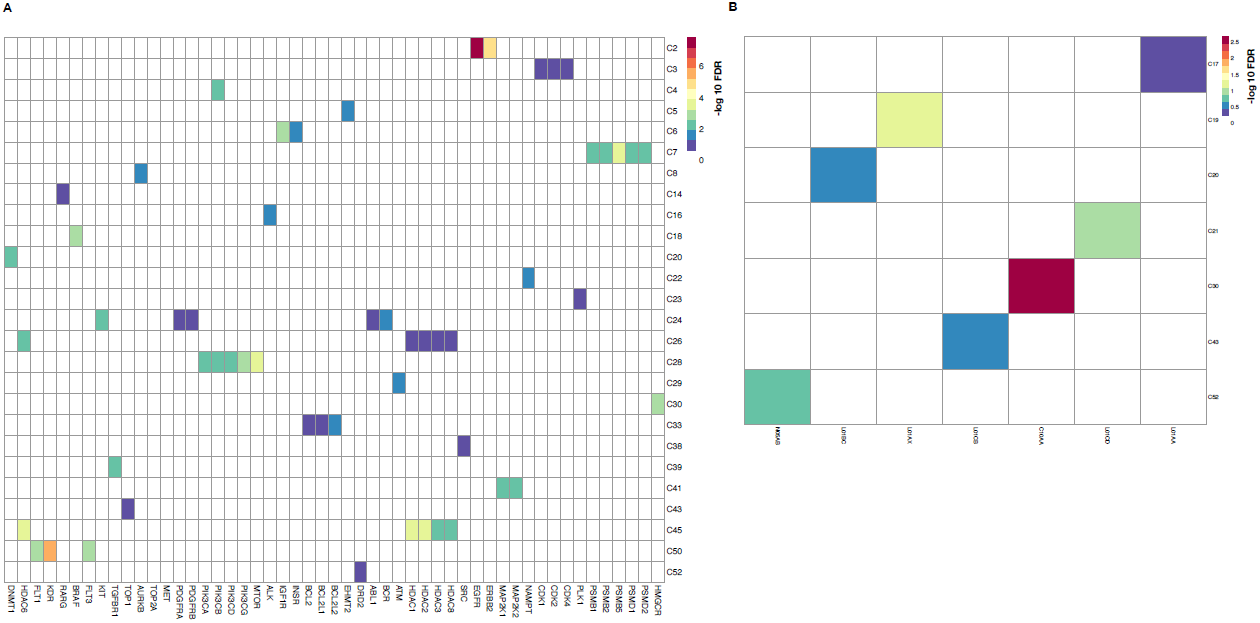
Enrichment of Drug Communities of the DNF taxonomy (using CTRPv2 sensitivity data). A total of 53 communities were tested for enrichment against drug target annotations from the CTRPv2 data and ATC annotations from ChEMBL (see methods). A Fisher test was performed between all the drugs in each community versus all drugs attributed to a specific drug target or ATC class, and corrected for multiple testing (FDR correction). **(A)** Enrichment of communities for Drug target annotations, with -log 10 FDR values indicated in the heat map, which has been reduced to show significantly enriched communities. Communities are labelled by community number as determined by the affinity propagation clustering algorithm. **(B)** Enrichment of communities for ATC classes, with -log10 FDR values indicated in the heat map, which has been reduced to show significantly enriched communities. Communities are labelled by community number as determined by the APC algorithm.

Clustering of the DNF drug taxonomies identified a wide range of community sizes **(Supplementary Fig. 7)**, with a median community size of 4 drugs both CTRPv2 and NCI60-based DNF taxonomies. Many of these communities were significantly enriched for drug targets and ATC classes (Fig. 5, Supplementary Fig. 6). Among these, for example, community C2 in CTRPv2 is statistically enriched against the ERBB2 and EGFR targets and contains well known inhibitors for these targets, such as afatinib, erlotinib and lapatinib (Fig. 4, Fig. 5A). Similarly, C30 hosts almost all of the members of the statin family, which are known to affect the mevalonate pathway and HMGCR (Fig. 4, Fig. 5A). We identified enrichment of DNF communities (based on NCI60) against several representative ATC categories (Supplementary Fig. 6B). These include communities enriched for known anticancer and other therapeutic classes, including antimalarial drugs (C4, ATC P01BE [Artemisinin and derivatives], anthracyclines (C12, L01DB [anthracyclines and related substances]), antimetabolites (C9,ATC L01BB [purine analogues]), cholesterol lowering agents (C33, ATC C10AA [HMG CoA reductase inhibitors]), corticosteroids (C18, overrepresented in many ATC categories since they are indicated for a large number of medical conditions), vinca and taxanes alcalolds (C38 and C49; L01CD [taxanes] and L01CA [Vinca alkaloids and analogues], respectively) and protein kinase inhibitors (C20, L01XE). As expected, communities containing very few annotated targets or ATC classes do not demonstrate significant enrichment (**Supplementary Tables 11 and 12** for CTRPv2 and NCI60, respectively).

### Identification of novel drug-drug relationships and drug action mechanisms

We conducted an explorative analysis to identify potential mechanisms for existing drugs and for poorly characterized drugs in the set of drugs constituting the DNF network. We highlight here a number of examples for newly identified drug mechanisms against a variety of compounds. We identified a community of HMG Co-A reductase inhibitors (statins) composed of fluvastatln, lovastatln, and simvastatin (C30; Fig. 4). These are a class of cholesterol-lowering drugs, and which have been found to reduce cardiovascular disease. Interestingly, parthenolide clusters with this community, and has been experimentally observed to inhibit the NF-Kb inflammatory pathway in atherosclerosis and in colon cancer ^30,31^, thereby suggesting similar behavior to statin compounds. We also classified correctly drugs with unannotated mechanisms/targets in CTRPv2 such as ifosfamide, cyclophosphamide and procarbazine (C17; Fig. 4) which are known alkylating agents (ATC code: L01A). Furthermore, this was also true for docetaxel and paclitaxel (C21; Fig. 4), two taxanes drugs with unannotated target in CTRPv2 although known as sharing similar MoA (ATC code: L01CD).

Our integrative drug taxonomy was also able to identify targets for drugs with poorly understood mechanisms and to infer new mechanism for other drugs. Community C15, for example, contains tigecycline and Col-3 (Fig. 4); both are derivatives of the antibiotic tetracycline ^32^. Tigecycline is an approved drug, however its target is not characterized in humans. Col-3 showed antitumor activities by inhibiting matrix metalloproteinase ^32^. Interestingly, tosedostat (CHR-2797), a metalloenzyme inhibitor with antiproliferative potential 33,is also a member of this community. Another drug in this community, phloretin, is a natural compound with uncharacterized targets and has been recently shown to deregulate matrix metalloproteinases at both gene and protein levels ^34^. Our results suggest that matrix metalloproteinases would be the preferred target for drugs in this community. DNF also consolidated previous findings for drugs that may serve as tubulin polymerization disruptors, and which have not been previously classified as such. We identified a community of three drugs (C49; Fig. 4) in which LY2183240, and YK-4-279 have been recently identified to decrease alpha-tubulln levels ^21^. Tivantinib, a c-MET tyrosine kinase inhibitor, also blocked microtubule polymerization ^35^. Interestingly, this community is tightly connected to known microtubule perturbagens (C21; Fig. 4).

The identification of community C33 including the BCL-2 inhibitors ABT-737 and navitoclax (Fig. 4) concurs with the study of Rees et al. ^36^ where a high expression of BCL-2 was reported to confer sensitivity to these two drugs. This was not the case for another BCL-2 inhibitor, obatoclax, for which they proposed that its metabolic modification in cells impacts its interaction with BCL-2 proteins, therefore reducing its potency. We showed indeed that obatoclax did not cluster with the other two BCL-2 inhibitors (ABT-737 and navitoclax). Such an example demonstrates how the structural and sensitivity profiles of these two BCL-2 inhibitors are largely coherent, as opposed to obatoclax, which previously showed off-target effects compared to ABT-737 ^37^. This provides a good evidence to consider sensitivity profiles when developing new potent and specific BCL-2 inhibitors.

Our results also suggest the existence of "super communities”, that are a grouping of several communities contributing to a larger, systems-based MoA. An example is provided by the tightly connected communities C3, C21, C23, C43 (Fig. 4). One of these communities (C3: Alvocidib, PHA-793887 and staurosporine) includes well-characterized inhibitors of cyclin dependant kinases (CDKs) that are known to be major regulators of the cell cycle. BMS-345541 for example, which also clusters with drugs in C3, is an ATP non-competitive allosteric inhibitor of CDK ^38^. Those compounds are positioned close in the community network to topoisomerase I and II inhibitors (C43: SN-38, topotecan, etoposlde, tenlposlde), mlcrotubule dynamics perturbators (C21: paclitaxel, docetaxel, vincristine, parbendazole) and polo-like kinase inhibitors (C23: GSK461364, GW843682X). Iorio *et al.* reported that the similarity between CDK inhibitors and the other DNA-damaging agents is mediated through a p21 induction, which explains the interconnection and rationale of similar transcriptional and sensitivity effects of these regulators of cell cycle progression ^12^.

DNF based on NCI60 sensitivity information enabled identification of three tightly connected drug communities: C2, C5, C32 **(Supplementary Fig. 5)**. These communities contain a number of compounds which showed antitumor activity by generating reactive oxygen species (ROS) (communities C2: elesclomol, fenretinide; C5: ethacrynic acid, curcumin; C32: bortezomib, menadione). Interestingly, ethacrynic acid, an FDA approved drug indicated for hypertension, clustered with curcumin, a component of turmeric. Ethacrynic acid inhibits glutathione S-transferase (GSTP1) and induced mitochondrial dependant apoptosis through generation of ROS and induction of caspases ^39^. Curcumln showed antltumor activity by production of ROS and promotion of apoptotic signaling. Thus, our results suggest that GSTP1 could be a potential target of the widely-used natural compound curcumin. Interestingly, some of the identified communities using NCI60, such as the tight connection between BRAF/MEK inhibitor drugs (C42; Supplementary Fig. 5), had also been identified in our original analysis using CTRPv2 sensitivity profiles. This demonstrates a high degree of specificity of drug-target associations across cell lines and experimental platforms, which is crucial in biomarker identification and translational research.

## DISCUSSION

Identification of drug mechanisms for uncharacterized compounds is instrumental for determination of on-targets responsible of pharmacological effects, and off-targets associated with unexpected physiological effects. Shortcomings of current approaches include a degree of reliance on pharmacological, biochemical, and functional annotations that pertain to existing, well-characterized drugs, and which may not be applicable towards prediction of a new small compounds ^15^,^40^. To address this issue, we developed DNF, a high-throughput, systematic and unbiased approach that relies on basic and complementary drug characteristics, and harnesses this integrative classification to provide a global picture of drug relationships.

In this analysis, we have conducted to our knowledge the first large-scale integration of high-throughput, drug-related data that encompasses drug structure, sensitivity and perturbation signatures towards elucidating drug-drug relationships. We have removed any reliance on existing annotations that pertain to existing drugs, such as drug-target classifications or knowledge of the anatomical and organ system targeted by the drug compounds. As a consequence, we developed a scalable approach that relies only on basic drug information, making DNF both flexible for comprehensive drug classification, but also adaptable for testing new experimental compounds with minimal information (Fig. 6).

**Figure 6.**
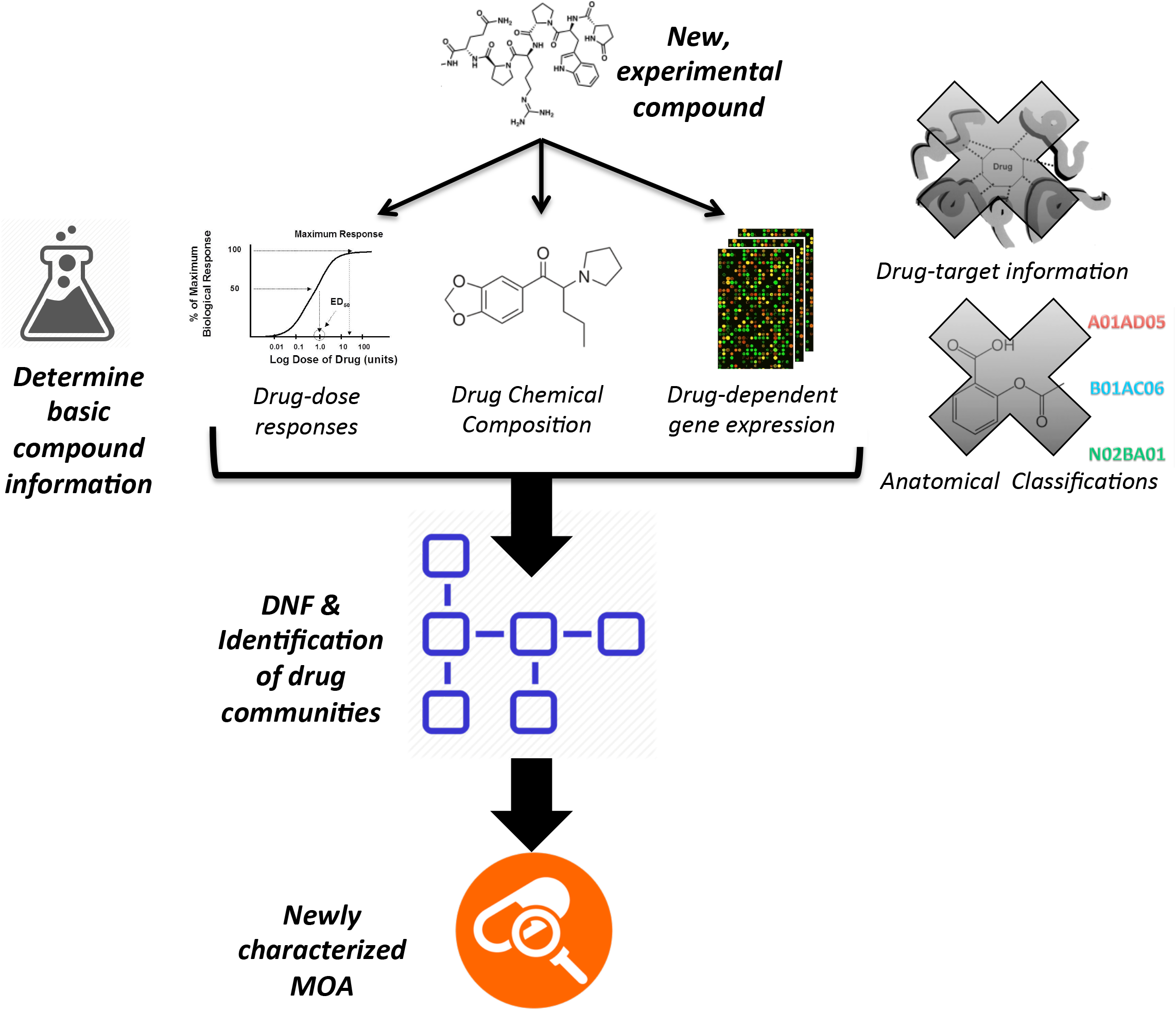
Schematic of the adaptability of DNF towards prediction of new experimental compounds.

Our high-throughput drug similarity network (DNF) capitalized upon our integrative Similarity Network Fusion method ^22^ to construct a global drug taxonomy based on the fusion of drug structure, sensitivity, and perturbation data. The construction of drug-similarity networks and their subsequent fusion allows us to harness the complementary nature of several drug datasets to infer an integrative drug taxonomy. Testing how well different drug taxonomies can correctly predict drug targets (Fig. 3A-B) and anatomical (ATC) drug classifications (Fig. 3C-D) indicates that DNF constitutes a significant improvement towards drug classification, compared to single data type analyses using either drug sensitivity, structure, or perturbation information alone. The marked improvement of drug classification using the similarity network-based method is sustained even with the use of a different source of *in vitro* sensitivity data (NCI60; Supplementary Fig. 4) to generate the DNF similarity matrix. Indeed, testing DNF using the NCI60 sensitivity information reveals that our integrative taxonomy consistently supersedes single-layer drug taxonomies across the target and ATC benchmarks despite the reduced number of cell lines used for sensitivity screening **(Supplementary Fig. 4)**. While DNF was not intended as a supervised approach to predict drug targets and ATC classifications *per se,* the ability to efficiently predict different drug classes provide credence to using our novel similarity network-based method to discover drug relationships. DNF is the only method that is consistently the top performer, while each single layer taxonomy performs well in only a few cases (Fig. 3 and Supplementary Fig. 4). Overall, these observations indicate that our integrative approach succeeds in combining several drug data types into a single comprehensive network that efficiently leverages the spectrum of the underlying data.

Our explorative analysis stresses the importance of drug sensitivity information as an important asset for prediction of drug-target associations (Fig. 3A,B and Supplementary Fig. 4A,B). Such findings support the relevance of pharmacological assays to predict drug targets, and underscore the comprehensive nature of the CTRPv2 dataset (860 cell lines screened with 16 drug concentrations, tested in duplicate) ^21^. Similarity, we have observed a priority for drug structure information towards prediction of ATC drug classification (Fig. 3C,D, Supplementary Fig. 4C,D). We have also conducted a quantitative comparison of the predictive performance of DNF against four published drug classification algorithms ^5,9,10,41^ that could be directly compared to the DNF approach (Supplementary Methods, Supplementary Fig. 8, **Supplementary Tables 13-18**). DNF outperforms the published methods in all cases **(Supplementary Fig. 8)**.

In addition to classification of drugs based on their targets and functional annotations, we also conducted an independent clustering of all of the drugs in the DNF network, and highlighted many cases of drug clusters (drug communities) with known MoA, thereby capturing context-specific features associated to drug sensitivity and transcriptomic profiles in cancer cells (Fig. 4, Supplementary Fig. 5). These cases serve both as ‘positive controls’ as well as validation of our methods. We demonstrated that DNF correctly identified communities of BRAF inhibitors, and MAPK/MEK inhibitors, among others in the CTRPv2 taxonomy (Fig. 4). We additionally highlight several well-characterized drug communities using the DNF taxonomy based on the NCI60 dataset **(Supplementary Fig. 5)**. Our quantitative assessment of the clusters identified from the DNF network revealed several communities that were significantly enriched for drug targets as well as ATC classes (Fig. 5, Supplementary Fig. 6), which underscores the biological relevance of the drug clusters that we had identified. Collectively, these findings support the use of DNF for the classification of drug relationships across several classes of drugs. Importantly, we compared the DNF drug communities to previously published data, and found an overlap between our results and clinical observations. For example, ibrutinib, which is a Bruton tyrosine kinase inhibitor (BTK) approved for the treatment of Mantle cell lymphoma and chronic lymphocytic leukemia, clustered with the known EGFR inhibitors (C2: erlotlnlb, gefltlnlb, afatinib and others). The effect of ibrutinib in EGFR Mutant Non-Small Cell Lung Cancer has been reported in a recent clinical trial (CllnlcalTrlals.gov Identifier: NCT02321540). This was also the case for MGCD265, a Met inhibitor, which clustered with most of the VEGFR (vascular endothelial growth factor receptor) inhibitors (C50: pazopanib, cediranib and others). In this community, pazopanib is the only FDA approved drug for the treatment of renal cell carcinoma. There exist a recent evidence that the clinical drug candidate MGCD265 has an application in renal malignancies (CllnlcalTrlals.gov Identifier: NCT00697632).

The current availability of sensitivity data and its overlap with drug perturbation and drug structure information remains a limiting factor. In our study we used the LINCS L1000 ^42^ dataset for drug perturbation because it contains much more drugs than the previous CMAP ^11^ (L1000=20,326 vs CMAP=1,309). Similarly we used sensitivity profiles from the CTRPv2 ^21^ and NCI60 datasets because they contain a large number of drugs compared to CCLE ^43^ or other smaller datasets (CTRPv2=481 and NCI60=49,938 vs CCLE=24). Our taxonomy is currently composed of nearly 240 drugs (using either CTRPv2 or NCI60 sensitivity data). The number of drugs with multiple data layer is likely to increase in the close future, as the LINCS L1000 is also expanding at fast pace and new sensitivity data are frequently released ^44^,^45^. Therefore, we developed DNF with scalability in mind as we expect the number of drugs in our network to increase continuously over time. Recognizing that the exploration of large-scale drug similarity network is challenging, we developed the DNF web-application to interactively display the CTRPv2 and NCI60 drug similarity networks (dnf.pmgenomlcs.ca). We leveraged the cytoscape.js library ^46^ to allow users to easily navigate drug communities and investigate the drugs within each community and their similarities.

The recent release of large-scale pharmacogenomic datasets, such as those generated within the CTRPv2 and LINCS L1000 projects, provides a unique opportunity to further investigate the effects of approved and experimental drugs on cancer models and their potential mode of action. Accordingly, future analyses for in-depth analysis of the molecular profiles of cancer cell lines, including mutation, CNV, transcription, methylatlon and proteomlc data, can reveal new biological processes associated with drug MoA. As a proof-of-concept, we investigated the molecular features associated with drug response in community C2 (*n*=9 drugs)from the CTRPv2 taxonomy (Fig. 4). We tested to which extent, ERBB2/EGFR pathways are correlated with the corresponding drug sensitivity from CTRPv2. We extracted basal gene expression profiles of the CTRPv2 cell lines (the same cell lines were profiled by the CCLE study), and correlated the gene expression profiles of cancer cell lines with the corresponding AUC values extracted from PharmacoGx ^23^. Single sample gene set enrichment analysis (ssGSEA) ^47^ was performed to identify pathways enriched in genes associated to sensitivity to the drugs in community C2, and ranked them with respect to their median enrichment score. Our results indicate that the ERBB2/EGFR pathway (PID_ERBB_NETWORK_PATHWAY from MSigDB ^48^) is highly correlated with sensitivity to the drugs in community C2 **(Supplementary Fig. 9)**. This was expected as most of them are known EGFR/ERBB2 inhibitors. Interestlngly,Ibrutlnlb, a BTKi (Bruton kinase inhibitor) showed a similar profile as EGFR inhibitors ^49^. This example shows how DNF can be used to investigate the molecular basis of the MoA shared by multiple drugs within a community. Methods leveraging DNF in combination with the wealth of molecular profiles from cancer cell lines and patient tumors holds the potential to to identify robust biomarkers and improve drug matching for individual patients, which would constitute a major step towards precision medicine and drug repurposing in cancer.

This study has several potential limitations. First, the number of drugs with all data layers available is limited by the small overlap between drug sensitivity and perturbation datasets. As these datasets grow, the DNF taxonomy will expand proportionally. Second, the L1000 is a relatively new dataset and there is no consensus yet on how to best normalize the data and compute the transcriptomic perturbation signature for each drug. In this study, we used the gene expression data as normalized by the Broad Institute (QNORM, level 3 data) and the signature model implemented in PharmacoGx ^23^; however we recognize that other pipelines could be used to mine the L1000 dataset and potentially improve DNF. Third, the low number of cell lines in specific tissue types ln L1000 prevented us from creating tissue-specific integrative taxonomies to better explore the molecular context of drug MoA. This limitation could be lifted with the availability of perturbation profiles for more cell lines in the future. Fourth, given that SuperPred and DrugE-Rank websites only report the top predictions but not the full list of drugs with similar targets or ATC codes, we computed the distance between these partial rankings to compute similarities between drugs (see Methods). Similarly, a direct comparison between DNF and the taxonomy inferred using Iorio *et al.* and Iskar *et al.* methods is not possible due their reliance on CMAP. The adaptation of these methods for the L1000 dataset was challenging due to the reduced set of 1000 genes. Consequently, the results of the comparison between DNF and published methods should be interpreted cautiously. Finally, we had to use different drug sensitivity measures for CTRPv2 and NCI60 as both projects released different types of pharmacological profiles: CTRPv2 reported percentage of cell viability while NCI60 reported percentage of growth inhibition controlled for population doubling time. Despite these heterogeneous drugs sensitivity data, we observed similar communities for the drugs in common between the two sensitivity datasets, supporting the robustness of DNF.

In conclusion, we developed an integrative taxonomy inference approach, DNF, leveraging the largest quantitative compendiums of structural information, pharmacological phenotypes and transcriptional perturbation profiles to date. We used DNF to conduct a cross¬comparative assessment between our integrative taxonomy, and single-layer drug taxonomies,as well as published methods used to predict drug targets or functional annotations. Our results support the superiority of DNF towards drug classification, and also highlights singular data types that are pivotal towards prediction of drug categories in terms of anatomical classification as well as drug-target relationships. Overall, the DNF taxonomy has produced the best performance, most consistent classification of drugs for multiple functional classes in both CTRPv2 and NCI60. The comprehensive picture of drug-drug relationships produced by DNF has also succeeded in predicting new drug MoA. The integrative DNF taxonomy has the potential to serve as a solid framework for future studies involving inference of MoA of new, uncharacterized compounds, which represents a major challenge in drug development for precision medicine.

## MATERIAL AND METHODS

A schematic overview of the analysis design is presented in Supplementary Fig. 1.

### Processing of drug-related data and identification of drug similarity

***Drug structure annotations:*** Canonical SMILES strings for the small molecules were extracted from PubChem ^50^, a database of more than 60 millions unique structures. Tanimoto similarity measures ^51^ between drugs were calculated by first parsing annotated SMILES strings for existing drugs through the *parse.smiles* function of the *rcdk* package (version 3.3.2). Extended connectivity fingerprints (hash-based fingerprints, default length 1,024) across all drugs was subsequently calculated using the *rcdk::get.fingerprints* function ^52^.

***Drug perturbation signatures:*** We obtained transcriptional profiles of cancer lines treated with drugs from the L1000 dataset recently released by the Broad Institute ^42^, which contains over 1.8 million gene expression profiles of 1000 ‘landmark’ genes across 20,413 drugs. We used our PharmacoGx package (version 1.3.4) ^23^ to compute signatures for the effect of drug concentration on the transcriptional state of a cell, using a linear regression model adjusted for treatment duration, cell line identity, and batch to identify the genes whose expression is significantly perturbed by drug treatment:

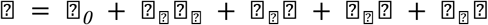

where

*G* = *molecular feature expression (gene)*
*C_j_* = *concentration of the compound applied*
*T* = *cell line identity*
*D* = *experiment duration*
*D* = *experimental batch*
*β* = *regression coefficients*.

The strength of the feature response is quantified by. and are scaled variables (standard deviation equals to 1) to estimate standardized coefficients from the linear model. The transcriptional changes induced by drugs on cancer cell lines are subsequently referred to throughout the text as *drug perturbation signatures.* Similarity between estimated standardized coefficients of drug perturbation signatures was computed using the Pearson correlation coefficient, with the assumption that drugs similarly perturbing the same set of genes might have similar mechanisms of action.

***Drug sensitivity profiles:*** We obtained summarized dose-response curves from the published drug sensitivity data of the NCI60 ^19^ and CTRPv2 ^21^ datasets integrated in the PharmacoGx package. We relied on the calculated Z-score and area under the curve (AUC) metrics from drug-dose response curves of the NCI60 and CTRPv2, respectively. Drug similarity was defined as the Pearson correlation of drug sensitivity profiles.

### Development of a drug network fusion (DNF) taxonomy

We used our Similarity Network Fusion algorithm ^22^ to identify drugs that have similar mechanisms of actions by integrating three data types representing drug structure, drug perturbation, and drug sensitivity profiles. Drug structure and drug perturbation taxonomies were based on drug-drug similarity matrices computed from the PubChem SMILES and the L1000 dataset, respectively. To test the robustness of the fusion algorithm with respect to the scale of the drug sensitivity profiles, we also applied our methodology on both the CTRPv2 and NCI60 datasets. The NCI60 dataset comprises a much smaller panel of cell lines (60 vs. 860 for NCI60 and CTRPv2, respectively). The NCI60 panel compensates for its small cell line panel by the large number of screened drugs (>40,000 drugs tested on the full panel; Supplementary Fig.2). Accordingly, the drug sensitivity taxonomy was composed of the drug-drug similarity matrix of the sensitivity profiles extracted from either the NCI60 or CTRPv2 datasets. For each dataset, an affinity matrix was first calculated using the *affinityMatrix* function as described in the *SNFtool* package (version 2.2), using default parameters. We combined the three affinity matrices of the structure, perturbation, and sensitivity taxonomies into a Drug Network Fusion (DNF) matrix using the *SNFtool::SNF* function **(Supplementary Fig. 1)**. Two separate DNF matrices were generated dependant on the sensitivity layer used (either CTRPv2 or NCI60). The developed DNF taxonomies, as well as the single data type taxonomies, were subsequently tested against benchmark datasets to validate their drug mode of action (MoA).

### Assessment of drug mode of action across drug taxonomies

***Drug-target associations.*** Known target associations for drugs pertaining to the NCI60 dataset were downloaded from DrugBank (version 5.0) ^53^. Drug-target associations for drugs of the CTRPv2 dataset were obtained from the CTRPv2 website (http://www.broadinstitute.org/ctrp.v2/?page=#ctd2Target). Drugs with annotated targets were filtered to retain only targets with at least two drugs.

*Anatomical therapeutic classification system (ATC).* ATC annotations ^54^ for the drugs common to the NCI60 and CTRPv2 datasets were downloaded from ChEMBL (file version 16-5¬10-02)^55^. These ATC codes were filtered to retain only those categories with at least one pair of drugs sharing a pharmacological indication. The drugs with known ATC annotations from the NCI60 and CTRPv2 datasets were subsequently used as a validation benchmark against singular drug taxonomies and the DNF taxonomy.

### Evaluation of drug mechanism of action across taxonomies

We assessed the predictive value of our developed taxonomies against drug-target and ATC benchmark datasets to determine the extent to which single data type taxonomies and the DNF taxonomy recapitulate known drug MoA **(Supplementary Fig. 3)**. We adapted the method from Cheng et al ^56^ to compare benchmarked datasets against singular drug taxonomies (drug perturbation, drug structure, or drug sensitivity) as well as the integrated DNF taxonomy. This method is further detailed below for the benchmark datasets used in our study. First, we created adjacency matrices that indicate whether each pair of drugs share a target molecule or ATC annotation. The drug-target and ATC adjacency matrices were then converted into a vector of similarities between every possible pair of drugs where the value ‘1’ was assigned in the vector if the paired drugs were observed the same target/ATC set, and ‘0’ otherwise. Similarly, the affinity matrices of singular drug taxonomies as well as the DNF taxonomy matrix were converted into vectors of drug pairs, with the similarity value of the drug pairs retained from their original corresponding matrix. Binary vectors of the benchmarks were compared to the four continuous vectors of the drug taxonomies by computing the receiver-operating curves (ROC) and the area under the curve (AUROC) using the *ROCR* package (version 1.0.7) ^57^. Concordance indices were calculated using the *rcorr.cens* function of the *Hmisc* package (version 1.18.0). The AUROC estimates the probability that, for two pairs of drugs, drugs that are part of the same drug set (same therapeutic targets or ATC functional annotations) have higher similarity than drugs that do not belong to the same drug set. AUROC calculations for each of the four taxonomies were statistically compared against each other using an adapted version of the *survcomp::compare.cindex* function ^58^. Precision-Recall (PR) curve is an alternative to ROC curves for measuring an algorithm’s performance, especially in classification problems with highly skewed class distributions ^59^. We used PRROC package (version 1.1) to compute PR curves. This package does not implement functions for statistical comparison of PR curves. For these types of classification tasks, algorithms that optimize AUROC do not necessarily optimize the area under the PR curves (AUPRC) ^59^. Therefore, computing both curves brings more insight to measuring performance and comparing multiple algorithms for the same prediction task.

### Detection of drug communities and visualization

Clusters of drug communities were determined from the DNF taxonomy using the affinity propagation algorithm ^27,60^ from the *apcluster* package (version 1.4.2). The apcluster algorithm generates non-redundant drug communities, with each community represented by an exemplar drug. An elevated *q* value parameter, which determines the quantiles of similarities to be used as input preference to generate small or large number of clusters, was set at *q*=0.9 within the *apcluster* function to produce a large number of communities. Networks of exemplar drugs were rendered in *Cytoscape* (version 3.3.0) ^61^. Drug structures were rendered using the *chemViz* plugin (version 1.0.3) for cytoscape ^62^. A minimal spanning tree of the exemplar drugs was determined using Kruskal’s algorithm as part of the *CySpanningTree* plugin (version 1.1) ^63^ for cytoscape.

### Assessment of Drug Community Enrichment

We tested for the enrichment of drug communities determined from the DNF taxonomy against all drugs attributed to a specific target or ATC class. We first generated lists of drug targets and ATC classes from our benchmarks, filtered to retain only drug targets or ATC classes with two or more drugs. Each of the 53 and 51 communities from the DNF taxonomy (using CTRPv2 and NCI60 drug sensitivity data, respectively) was subsequently compared against the lists using a Fisher test, followed by multiple testing (FDR) correction.

### DNF web-application

We developed the DNF web-application (dnf.pmgenomics.ca) to facilitate the exploration of the drug communities and the full drug similarity networks built using the CTRPv2 and NCI60 drug sensitivity datasets. The application was implemented using JavaScript and AngularJS ^64^ for its frontend and Node.js ^65^ for its backend. Drug network information is stored on the server as JSON and rendered as graphs with the Cytoscape.js library ^46^. As drug clusters are fully connected, showing all edges in the web application will overwhelm the browser. Thus, the graphs shown in the web application are thresholded to display the top one thousand edges between drugs that have the greatest similarity. Users can click on drug communities to display the full set of drugs and their similarity edges. Clicking on an edge reports the similarity score and its relative contribution to the fused similarity score. Clicking on drugs provides basic descriptors of the compound and its pubchem link.

### Research reproducibility

The code and data links required to reproduce this analysis is publicly available on github.com/bhklab/DNF. All software dependencies are available on the Comprehensive Repository R Archive Network (CRAN) ^66^ or Bioconductor (BioC) ^67^. A detailed tutorial describing how to run our analysis pipeline to generate the figures and tables are provided in Supplementary Information. The procedure to setup the software environment and run our analysis pipeline is also provided. This work complies with the guidelines proposed by Sandve et al. ^68^ in terms of code availability and replicability of results.

## ACKNOWLEDGEMENTS

The authors would like to thank Jacques Archambault for sharing his expertise in pharmacology and supporting Nehme El-Hachem financially, and Dr. Fei-Fei Liu for supporting Laleh Soltan Ghoraie financially. We thank John Chen and Ali Nawed for their contribution to the DNF web-application. They also thank Dr. Shanfeng Zhu for running the DrugE-Rank prediction algorithm on the set of drugs analyzed in this study.

### AUTHOR CONTRIBUTIONS

N.E-H, D.M.A.G and L.S.G contributed equally to this work. N.E.-H, D.M.A.G and L.S.G wrote the code, performed the analysis and interpreted the results. L.S.G compared DNF to other methods. Z.S and P.S collected and curated the drug perturbation and sensitivity datasets. A.G developed the SNF algorithm and assisted in its application to drug-related data. R.I. and G.D.B contributed to the analysis of drug communities and interpretation of the results. D.M.A.G, N.E-H,and B.H-K conceived the initial design of the study, L.S.G designed the comparison with competitive approaches from the literature. B.H-K supervised the study. D.M.A.G, N.E-H, L.S.G and B.H-K wrote the manuscript.

### FUNDING

This study was conducted with the support of the Canadian Cancer Research Society and the Ontario Institute for Cancer Research through funding provided by the Government of Ontario. Z.S was supported by the Cancer Research Society (Canada). N.E-H was supported by the Ministry of Economic Development, Employment and Infrastructure and the Ministry of Innovation of the Government of Ontario. B.H.K was supported by the Gattuso-Slaight Personalized Cancer Medicine Fund at Princess Margaret Cancer Centre and the Canadian Institutes of Health Research.

### COMPETING FINANCIAL INTERESTS

The authors declare no competing financial interests.

## SUPPLEMENTARY FIGURE LEGENDS

**Supplementary Figure 1:**
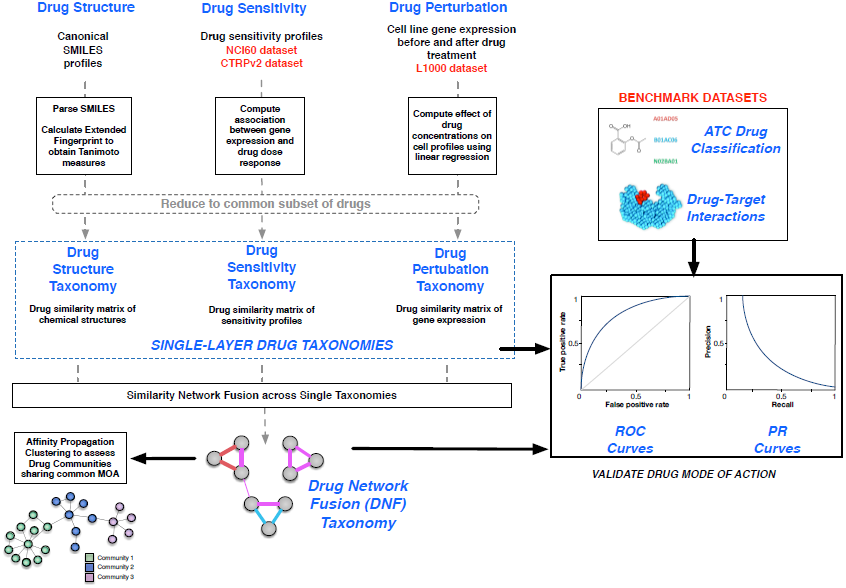
Overview of the study design. Drug sensitivity profiles from the NCI60 and the CTRPv2 datasets, along with drug perturbation and drug structure data from the L1000 dataset, are first parsed into drug-drug similarity matrices that represent single-dataset drug taxonomies. Two DNF taxonomies are generated using the drug sensitivity taxonomy from either the NCI60 or CTRPv2 datasets. DNF taxonomies and single-dataset taxonomies are tested against benchmarked datasets containing ATC drug classification and drug-target information, to validate their efficacy in predicting drug MoA. Additional clustering is conducted on DNF taxonomies to identify drug communities sharing a MoA.

**Supplementary Figure 2:**
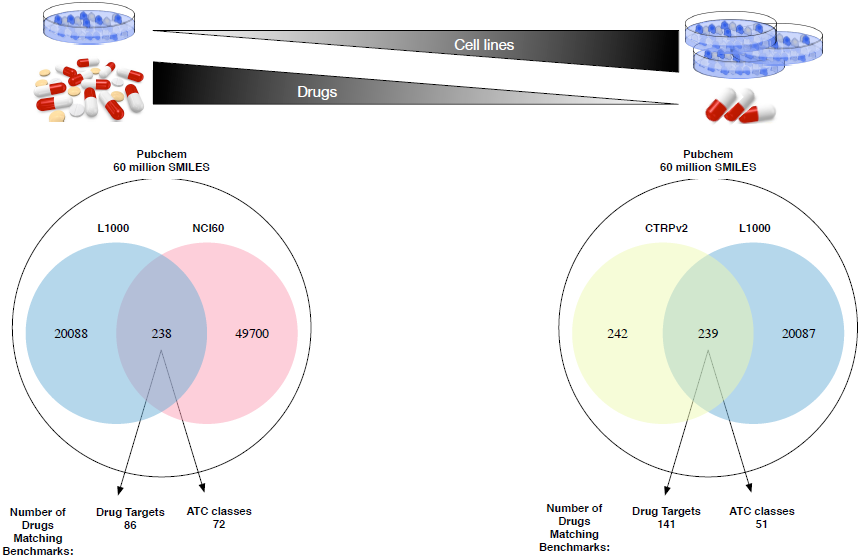
Overlap of drug annotations across the L1000 and the NCI60 and CTRPv2 sensitivity datasets. Also indicated are the number of drugs from each DNF matrix, which overlap with the drug target and ATC benchmarks.

**Supplementary Figure 3:**
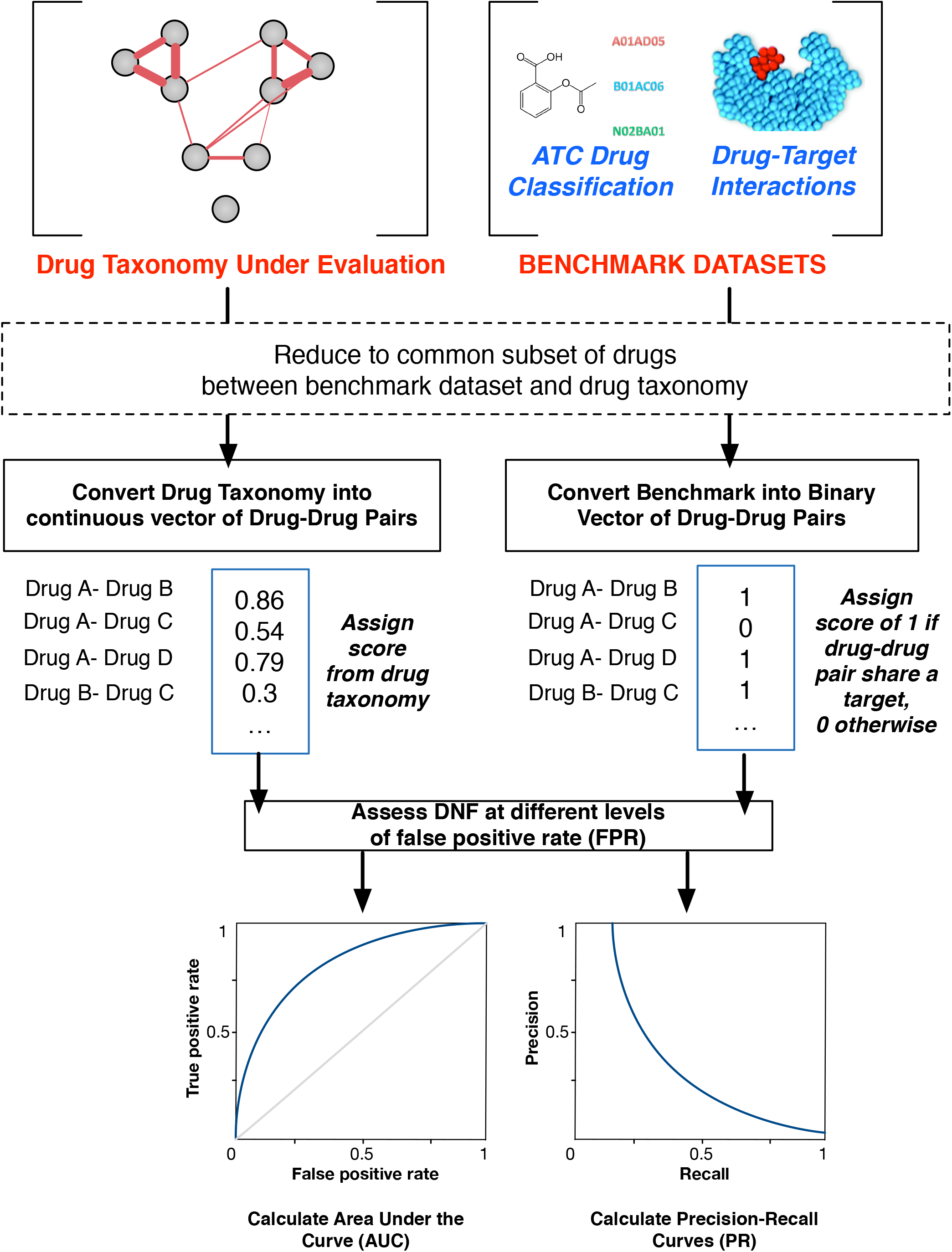
Schematic representation of the validation of the DNF and single data type analyses against drug benchmarks. Drug taxonomies are converted into a continuous vector of drug-drug pairs. Benchmark datasets are converted into binary vectors, whereby a given drug-drug pair is assigned a value of ‘1’ if the drugs share a common drug target or ATC classification, and ‘0’ otherwise. Vectors are compared using AUROC and AUPRC.

**Supplementary Figure 4:**
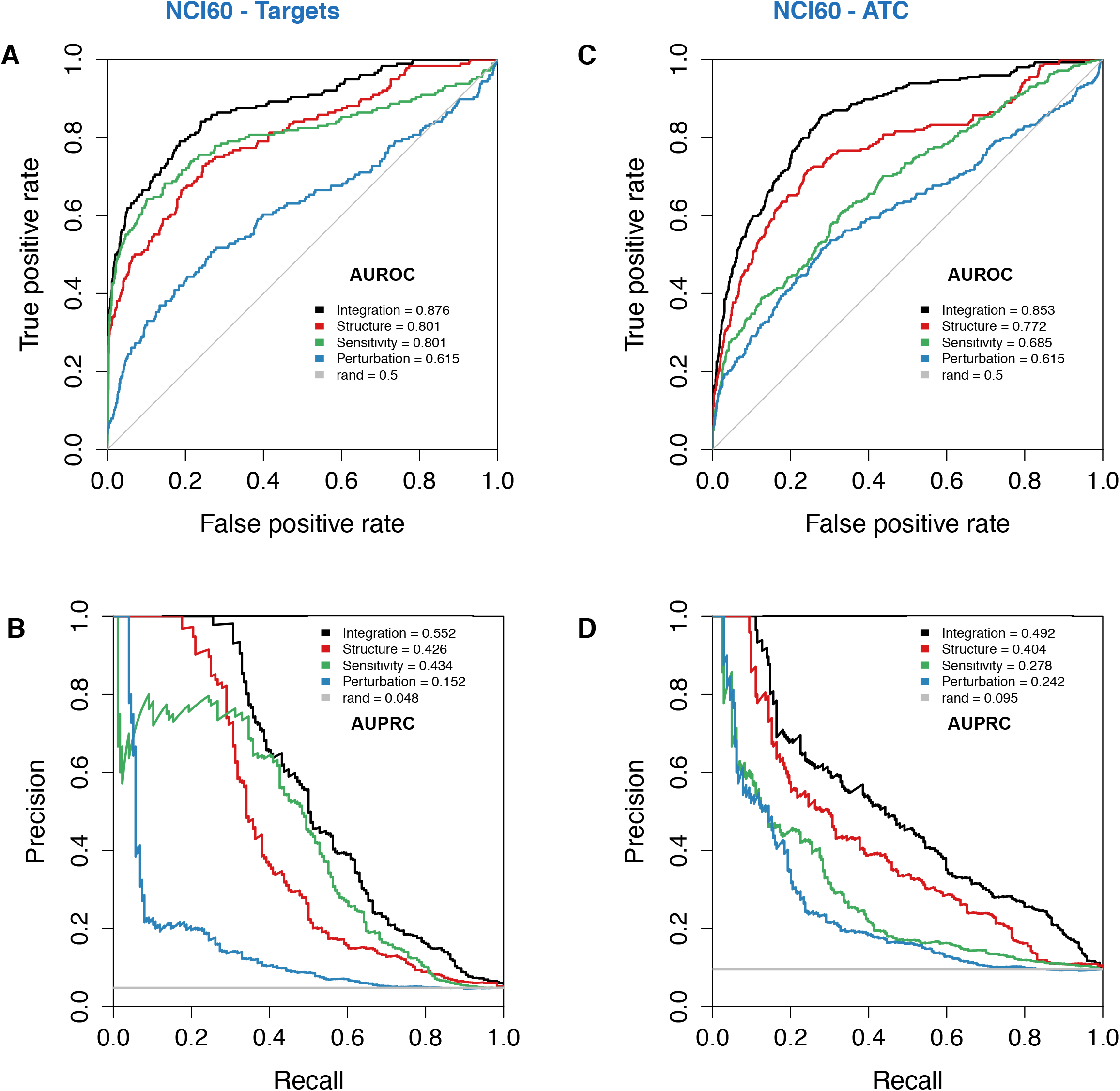
Validation of single-dataset and DNF taxonomies against drug benchmark datasets, based on DNF generated using NCI60. ROC and PR curves are shown for each of the taxonomies, tested against ATC annotations and drug-target information from Chembl or internal benchmarks. A diagonal (grey) representing the null case (AUROC=0.5) is drawn for clarity, and a grey line is also drawn to map random ‘rand’ cases for the PR curves. **(A)** ROC curve for NCI60 against drug-targets **(B)** PR curve against drug-targets **(C)** ROC curve for NCI60 against ATC **(D)** PR curve against ATC drug classifications.

**Supplementary Figure 5:**
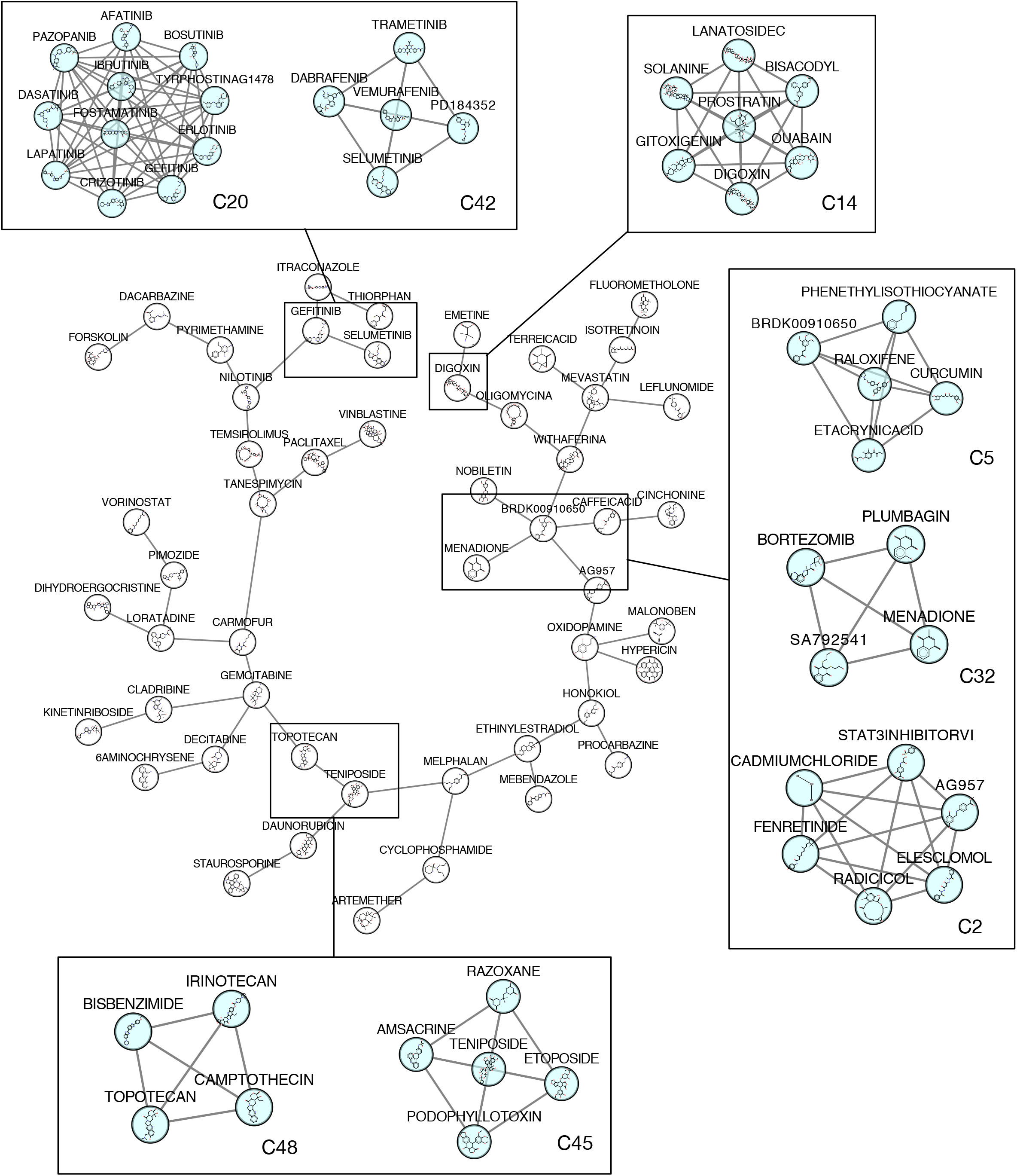
Community of 53 Exemplar drugs of the DNF taxonomy generated using NCI60. Communities sharing similar MoA and proximity in the network are highlighted, with the community number indicated.

**Supplementary Figure 6:**
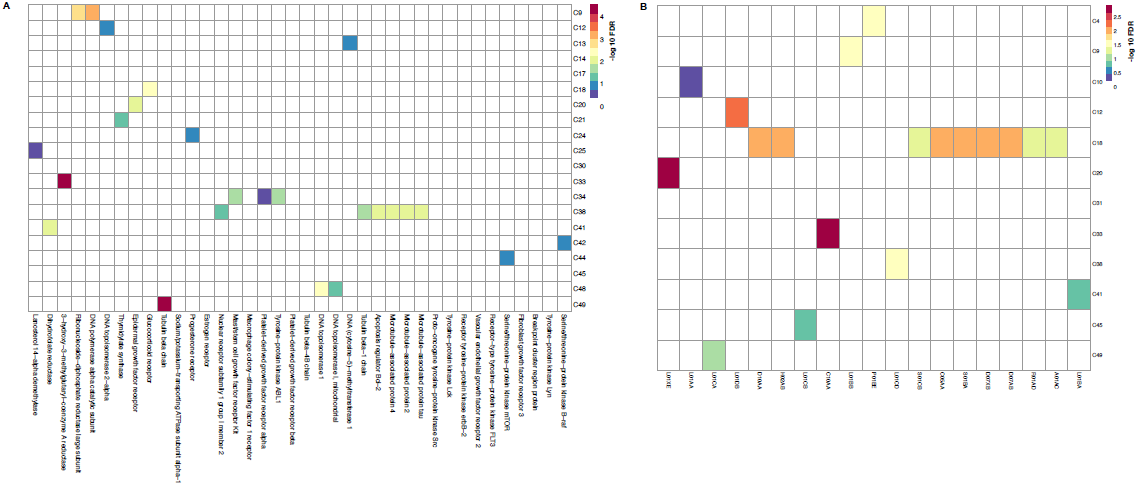
Enrichment of Drug Communities of the DNF taxonomy generated using NCI60. A total of 51 communities were tested for enrichment against drug target annotations from DrugBank and ATC annotations from ChEMBL. **(A)** Enrichment of communities for Drug target annotations, with -log10 values indicated in the heatmap, which has been reduced to show significantly enriched communities. Communities are labelled by community number as determined by the APC algorithm. **(B)** Enrichment of communities for ATC classes, with -log10 values indicated in the heat map, which has been reduced to show significantly enriched communities. Communities are labelled by community number as determined by the APC algorithm.

**Supplementary Figure 7:**
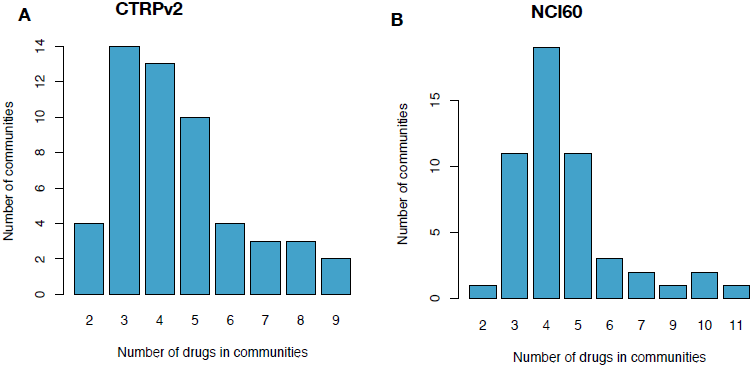
Distribution of drug communities sizes of the DNF taxonomy. **(A)** A total of 53 communities from DNF (using CTRPv2 sensitivity data) were tested for enrichment against drug target annotations from the CTRPv2 data and ATC annotations from ChEMBL (see methods). **(B)** A total of 51 communities from DNF (using NCI60 sensitivity data) were tested for enrichment against drug target annotations from DrugBank and ATC annotations from ChEMBL (see methods).

**Supplementary Figure 8:**
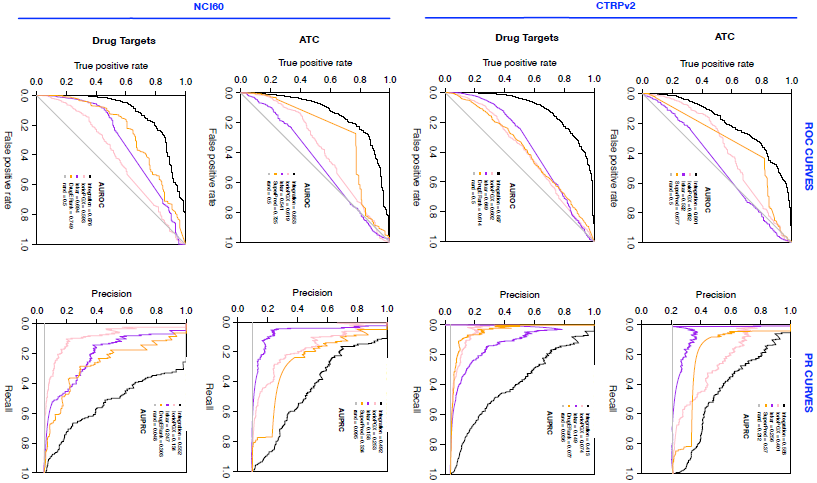
Comparison of the DNF and single-layer drug taxonomies with comparable drug prediction methods (SuperPred, DrugE-Rank, Iorio, Iskar). ROC and PR curves are depicted to indicate the performance of the taxonomies and comparative methods against drug target and ATC benchmarks. This analysis is conducted for both DNF taxonomies (based on CTRPv2 or NCI60 data, shown in blue), and their associated similarity networks. ROC curves are on the left, and PR curves on the right.

**Supplementary Figure 9:**
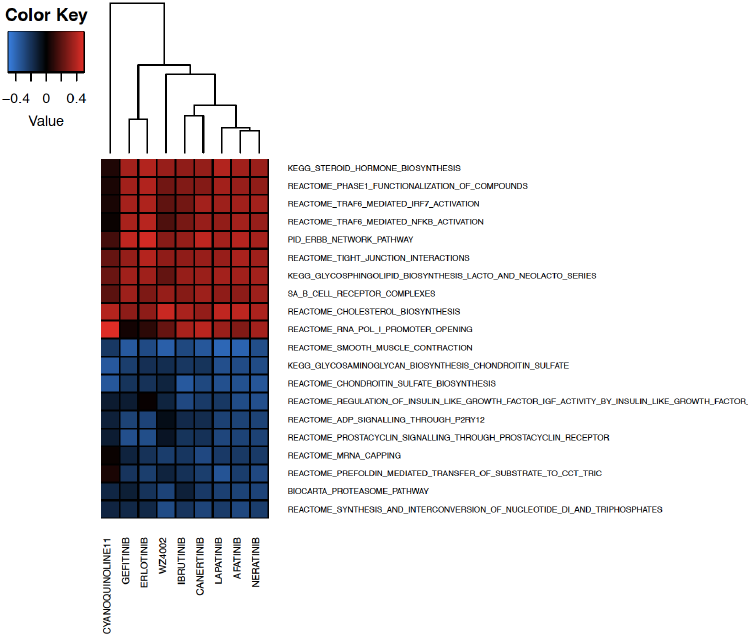
Assessment of Drug-pathway associations for community C2 of DNF (based on CTRPv2). We selected C2 (9 drugs) and tested to which extent, ERBB2/EGFR pathways are correlated with the corresponding drug sensitivity from CTRPv2. The top 10 up/downregulated pathways were shown in a heatmap.

## SUPPLEMENTARY TABLE LEGENDS

**Supplementary Table 1. Similarity Matrices of the DNF and single-layer taxonomies (based on the CTRPv2 drug sensitivity dataset)**. **(A)** Similarity Matrix of the fused DNF taxonomy **(B)** Similarity Matrix of the Perturbation layer **(C)** Similarity Matrix of the Sensitivity layer **(D)** Similarity Matrix of the Structure layer

**Supplementary Table 2. Similarity Matrices of the DNF and single-layer taxonomies (based on the NCI60 drug sensitivity dataset). (A)** Similarity Matrix of the fused DNF taxonomy **(B)** Similarity Matrix of the Perturbation layer **(C)** Similarity Matrix of the Sensitivity layer **(D)** Similarity Matrix of the Structure layer

**Supplementary Table 3. Matrix of Correlations between single-layer drug taxonomies, and correlations between DNF and single layers.** Data are shown for both **(A)** DNF using CTRPv2 and **(B)** DNF using the NCI60 sensitivity datasets.

**Supplementary Table 4. Statistical comparison of the DNF taxonomy against single datasets taxonomies, using one-sided superiority tests.** Comparisons were conducted for both DNFs generated using the CTRPv2 or the NCI60 datasets. Reported scores pertain to comparisons conducted using both drug benchmarks (Drug-target information as well as ATC).

**Supplementary Table 5. List of identified communities using the community detection algorithm against the DNF generated using CTRPv2.** Exemplar drugs for each community are identified, along with the number of drugs in that community. The list of drugs pertaining to each community is indicated. Drug populations are coloured to indicate communities that have in green means that they have at least 2 drugs with a known mechanism of action (total 139 drugs for CTRPv2, green), and those communities where drugs are unlabeled or unclassified (orange).

**Supplementary Table 6. Refined list of identified communities using the APC cluster algorithm against the DNF generated using CTRPv2, selected for communities that have at least two drugs with a known mechanism of action.** Exemplar drugs for each community are identified, along with the number of drugs in that community. The list of drugs pertaining to each community is indicated.

**Supplementary Table 7. Summary of Functional Drug Classes Identified Using DNF**

**Supplementary Table 8. Summary of communities generated from CTRPv2/L1000 integrative layers showing positive controls cases with at least 2 drugs sharing a mechanism of action from the same community.**

**Supplementary Table 9. List of identified communities using the community detection algorithm against the DNF generated using NCI60.** Exemplar drugs for each community are identified, along with the number of drugs in that community. The list of drugs pertaining to each community is indicated. Drug populations are coloured to indicate communities that have in green means that they have at least 2 drugs with a known mechanism of action, and those communities where drugs are unlabeled or unclassified (orange).

**Supplementary Table 10. Refined list of identified communities using the community detection algorithm against the DNF generated using NCI60, selected for communities that have at least two drugs with a known mechanism of action.** Exemplar drugs for each community are identified, along with the number of drugs in that community. The list of drugs pertaining to each community is indicated.

**Supplementary Table 11. List of enrichments of drug communities from DNF generated using CTRPv2 sensitivity against drug target (A) and ATC classes (B)**

**Supplementary Table 12. List of enrichments of drug communities from DNF generated using NCI60 sensitivity against drug target (A) and ATC classes (B)**

**Supplementary Table 13. Similarity matrix used in benchmarking of SuperPred (based on the CTRPv2 drug sensitivity dataset)**

**Supplementary Table 14. Similarity matrix used in benchmarking of SuperPred (based on the NCI60 drug sensitivity dataset)**

**Supplementary Table 15. Similarity matrix used in benchmarking of the Iskar algorithm**

**Supplementary Table 16. Similarity matrix used in benchmarking of the Iorio algorithm**

**Supplementary Table 17. Similarity matrix used in benchmarking of the Drug E-Rank algorithm (based on the CTRPv2 drug sensitivity dataset)**

**Supplementary Table 18. Similarity matrix used in benchmarking of the Drug E-Rank algorithm (based on the NCI60 drug sensitivity dataset)**

## SUPPLEMENTARY INFORMATION

## SUPPLEMENTARY METHODS

### Comparison of DNF with other methods

We have searched the literature for drug classification algorithms that are comparable with our integrative similarity-based network approach. Notably, many available methods incorporate ATC and drug target information as input variables for their predictions, which poses an obstacle for comparison, as DNF does not rely on these data types as input but uses them as external benchmarks. As such, we identified a limited number of state-of-the-art methods to perform a comparative study with DNF. These methods comprise two main groups. The first group attempt to decode drugs’ mechanism of action based on drug similarities from CMap perturbation data (Iorio et al. 2009; Iskar et al. 2010). The second group comprise mainly supervised machine learning methods applied for ATC or target prediction (Nickel et al. 2014; Yuan et al. 2016).

### Comparison to MoA decoding methods

Relying only on transcriptomic perturbation data, Iorio et al. and Iskar et al. applied different approaches to first, preprocess CMap’s perturbation profiles (Thiers 2007; Lamb et al. 2006) and then, to compute the same drug-drug similarity score. In order to calculate similarity between drugs *d_i_* and *d_j_* {j=1,..,*n*} (*n* is total number of drugs), first, a *signature* is defined for *d_i_*, that is, two sets of (*m*=) 250 most significantly up and 250 downregulated genes are selected from the perturbation profile of *d_i_*. Second, *connectivity scores* (Thiers 2007; Lamb et al. 2006), based on Gene Set Enrichment Analysis (GSEA), between *d_i_* and all *d_j_*s, are calculated and stored. The computed scores are not necessarily symmetric, i.e., score*_di,dj_* ≠ score*d_j_,d_i_*. The final score for each drug pair is calculated as the average of the two scores. Both lorio et al. and Iskar et al. used the final scores to construct a drug similarity network. We used the same approach to calculate the similarity scores for the set of drugs under study for DNF. However, the aforementioned studies rely on the CMap data, while we used the L1000 profiles in our study. Therefore, due to the smaller number of genes (978) in the L1000 dataset, we had to select a smaller size (e.g., *m*=30) for the drug signatures. Additionally, we had to adapt pre¬processing approaches used by the aforementioned studies to be applicable to L1000. The adaptations are described as follows.

### Pre-processing according to Iorio et al.

Following Iorio’s method, for each drug, we first aggregated all lists of differentially expressed genes computed from treating different cell lines by the drug. For this purpose, we used *RankMerging* function (*GeneExpressionSignature* package: http://www.bioconductor.org/packages/release/bioc/html/GeneExpressionSignature.html). The method uses computes Spearman’s foot-rule (distance measure between two ranked lists) between each pair of signatures, and using *Borda* merging method, repeatedly, it merges the most similar pair of ranked lists each time, till obtaining one single ranked list for the drug.

After aggregating the signatures, we calculated the drug-pair distances (*connectivity scores* described above). Unfortunately, the corresponding prediction results were not significant. Therefore, in the second attempt, we applied the pre-processed perturbation signatures by PharmacoGx (Smirnov et al. 2016) package. Then, we computed the scores, and calculated the prediction results (shown in **Supplementary Figure 8** as IorioPGX). The prediction results are close to the perturbation layer of DNF, as expected.

### Pre-processing according to Iskar et al.

Iskar et al. follow as very different approach from Iorio et al. for pre-processing drug signatures. In order to adapt the method to the L1000 dataset, we first, discarded vehicle controls and mean centered the treatment samples within each batch. Then, for each drug, we averaged all its treatment replicates for each cell line. Therefore, we ended up with one single list for each drug for each cell line. L1000 consists of 77 cell lines in contrast to only five cell lines in CMap. In the next step, we calculated *connectivity scores* between drugs within each cell line. For each drug pair, the scores over multiple cell lines were averaged to compute the final scores. Prediction results based on these scores have been demonstrated (**Supplementary Figure 8**).

### Comparison to supervised ATC/target prediction methods

The ATC (target) prediction methods decode ATC codes (targets) for unknown drugs according to their similarities to a set of drugs with known ATC codes (targets). We selected SuperPred (Nickel et al. 2014) and DrugE-Rank (Yuan et al. 2016) methods for comparison with DNF, as these methods are freely available. Therefore, these methods do not aim at providing drug similarity scores and therefore they are not directly comparable to DNF. To address this issue, we post-processed their outputs generated from the corresponding web-based applications.

### SuperPred: ATC prediction Method

SuperPred computes and integrates three structural, i.e., 2D, fragment and 3D, similarities between an input drug d_i_ and a dataset of drugs with known ATC codes.

The following steps were performed to collect SuperPred’s output and compare to DNF:

1. We retrieved SMILES for the set of drugs under study by DNF (d_i_) and submitted them one-by-one to the SuperPred website and obtained the predictions. SuperPred retrieves the top five similar drugs (d’_j_) to the input drug. A prediction score is assigned to each retrieved drug (score_di,d’j_). Each d’_j_ is a drug with a set of *k* known ATC codes (atc^k^_d’j_). In some cases, the top prediction (i.e., the most similar drug) retrieved by SuperPred is the same as the input drug. We discarded such predictions, and collected the rest of the predictions for each drug (see “superPredResultsFinal.xlsx”). We defined the set of ATC predictions for d_i_ based on ATCs of d’_j_s, i.e., ATC_di_ = {atc^k^_d‘j_}, and assigned a weight to each code according to the prediction score, i.e., w_atc,di_^k^ = score_di,d’j_.
2. We used Kendall's *tau* distance between partial rankings (Fagin et al. 2003) on SuperPred’s predictions to compute pairwise drug-drug similarities.
3. Finally, we processed this matrix using “generateDrugPairs.R”, “generateRocPlot.R” and “generatePRPlot.R” functions (https://github.com/bhklab/DNF) along with predictions from the four layers of DNF. Please refer to **Supplementary Figure 8** for results.

### DrugE-Rank: Target prediction Method

DrugE-Rank (Yuan et al. 2016), a state-of-the-art target prediction method that uses an ensemble of a few efficient computational methods, i.e., k-nearest neighbor (k-NN), Bipartite Local Model with support vector classification (BLM-svc) (Bleakley and Yamanishi 2009), Bipartite Local Model with support vector regression (BLM-svr) (Bleakley and Yamanishi 2009), Laplacian regularized least squares (LapRLS) (Xia et al. 2010), Network based Laplacian regularized least squares (NetLapRLS) (Xia et al. 2010), and Weighted Nearest Neighbor-based Gaussian Interaction Profile classifier (WNN-GIP) (van Laarhoven, Nabuurs, and Marchiori 2011) (van Laarhoven and Marchiori 2013). The predictions for our drug set were kindly provided by Dr. Shanfeng Zhu. For each drug, *d_i_* a ranked list of 20 targets was provided. Then, for each pair of drugs we followed steps 2 and 3 from the SuperPred’s post-processing algorithm, described above, to compute the pairwise similarities and to compare with DNF’s results.

### Cautionary Note

Although the adaptation of these methods was challenging due to the use of different perturbation data (L1000 instead of CMAP for lorio and Iskar) and the limitations of the SuperPred website (predictions restricted to the top five hits), we provided the results of our comparison in **Supplementary Figure 8**. While DNF outperforms the published methods in all cases, we acknowledge that these results should be cautiously interpreted due to the differences in both data pre-processing and modification of the algorithms to make them comparable with our similarity-based networks.

